# Improvements to Bayesian Gene Activity State Estimation from Genome-Wide Transcriptomics Data

**DOI:** 10.1101/241000

**Authors:** Craig Disselkoen, Nathan Hekman, Brian Gilbert, Sydney Benson, Matthew Anderson, Matt DeJongh, Aaron Best, Nathan Tintle

## Abstract

An important question in many biological applications, is to estimate or classify gene activity states (active or inactive) based on genome-wide transcriptomics data. Recently, we proposed a Bayesian method, titled *MultiMM*, which showed superior results compared to existing methods. In short, MultiMM performed better than existing methods on both simulated and real gene expression data, confirming well-known biological results and yielding better agreement with fluxomics data. Despite these promising results, *MultiMM* has numerous limitations. First, *MultiMM* leverages co-regulatory models to improve activity state estimates, but information about co-regulation is incorporated in a manner that assumes that networks are known with certainty. Second, *MultiMM* assumes that genes that change states in the dataset can be distinguished with certainty from those that remain in one state. Third, the model can be sensitive to extreme measures (outliers) of gene expression. In this manuscript, we propose a modified Bayesian approach, which addresses these three limitations by improving outlier handling and by explicitly modeling network and other uncertainty yielding improved gene activity state estimates when compared to *MultiMM*.

## I. Introduction

Many approaches to analyzing genome-wide transcriptomics data attempt to leverage the data by classifying genes into one of two *gene activity states: active* (roughly speaking, the gene product is part of an active cellular mechanism) or *inactive* (the cellular mechanism is not active) [1]–[3]. Previous methods were limited by (a) assuming similar activity state expression thresholds across genes, such as in GIMME [4] where a user-specified expression level is used to classify gene activity states in any given experiment, (b) assuming similar proportions of active genes across experiments/conditions, (c) ignoring *a priori* information about potential gene co-regulation and (d) failing to adequately incorporate statistical uncertainty in subsequent inference about gene activity states.

Recently, we published a novel approach using a Bayesian Gaussian mixture model, *MultiMM* [5]. Grounded in a rigorous statistical framework, *MultiMM* addresses these limitations as demonstrated by better performance than existing methods on simulated and real transcriptomics data and higher consistency with accepted biological results and fluxomics data [4].The *MultiMM* algorithm takes as input a genome-wide matrix of transcriptomics data *E* across many experimental conditions and estimates the gene activity state of each gene *i* in condition *j*. *MultiMM* allows for *a priori* specification of sets of genes which are known to be co-regulated so that they may be classified as all active or inactive in the same experimental condition. Unlike previous methods, the estimated mixture distribution parameters can be used to yield a posterior probability *a_ij_* ∊ [0,1], that gene *i* is active in condition *j*. Recently, we further demonstrated that use of gene activity estimates outperformed the use of raw expression data when conducting gene-set analysis approaches to test for differential gene expression [6]. The promising results of the *MultiMM* method, however, are tempered somewhat by at least three significant limitations. First, the *MultiMM* method assumed that *a priori* identification of co-regulated genes was certain. This is rarely the case. Depending on data quality, *a priori* sets of co-regulated genes are likely often a mix of genes, which are co-regulated, are co-regulated only in some conditions or are not co-regulated at all. Second, in the first step of the method, inference about whether a gene is changing states (mathematically, whether expression values come from a one- or two-component mixture model) is taken to be certain, but in reality these classifications are estimates. Finally, inference about whether a gene’s expression values are from one or two components, as well as the Gibbs sampling method to estimate the model’s parameter values, can be sensitive to extreme values (outliers). In this paper we present enhancements to the *MultiMM* approach which address the noted shortcomings of the existing method and evaluate the modified method compared to the existing *MultiMM* method.

## II. Methods

### A. Modifications to the existing MultiMM Approach

The following three sections discuss limitations and modifications to the *MultiMM* approach.

#### i. Hedging to improve inference about whether a gene is changing activity states

The current *MultiMM* method [5] uses the Bayesian Information Criterion (BIC) to assess the fit of one- or two-component Gaussian mixture models in order to determine if evidence exists that a gene (or co-regulated set of *p* genes) is not changing activity states across the set of experiments being evaluated (in which case there is only one component). Following Raftery [7], the *MultiMM* method assumes that the 2-component distribution is assumed to be the better fit unless the 1-component model has at least a 12-point lower BIC. Genes for which the 2-component distribution is a better fit are classified as “changing state,” and genes for which the 1-component distribution is a statistically significantly better fit are classified as “not changing state.” In some cases, the resulting activity state estimates vary dramatically depending on whether a gene is classified changing-state or not (see, for example, Figure 1). We would like to incorporate the uncertainty of whether or not a gene is changing state in downstream analyses. To do so, we introduce a modification to the algorithm that both (a) provides a smoother transition between the two approaches rather than a sharp cutoff; and (b) “hedges” its estimates in cases of uncertain classifications, providing more balanced predictions and more accurately incorporating uncertainty in downstream parts of the *MultiMM* procedure.

**Figure 1.**
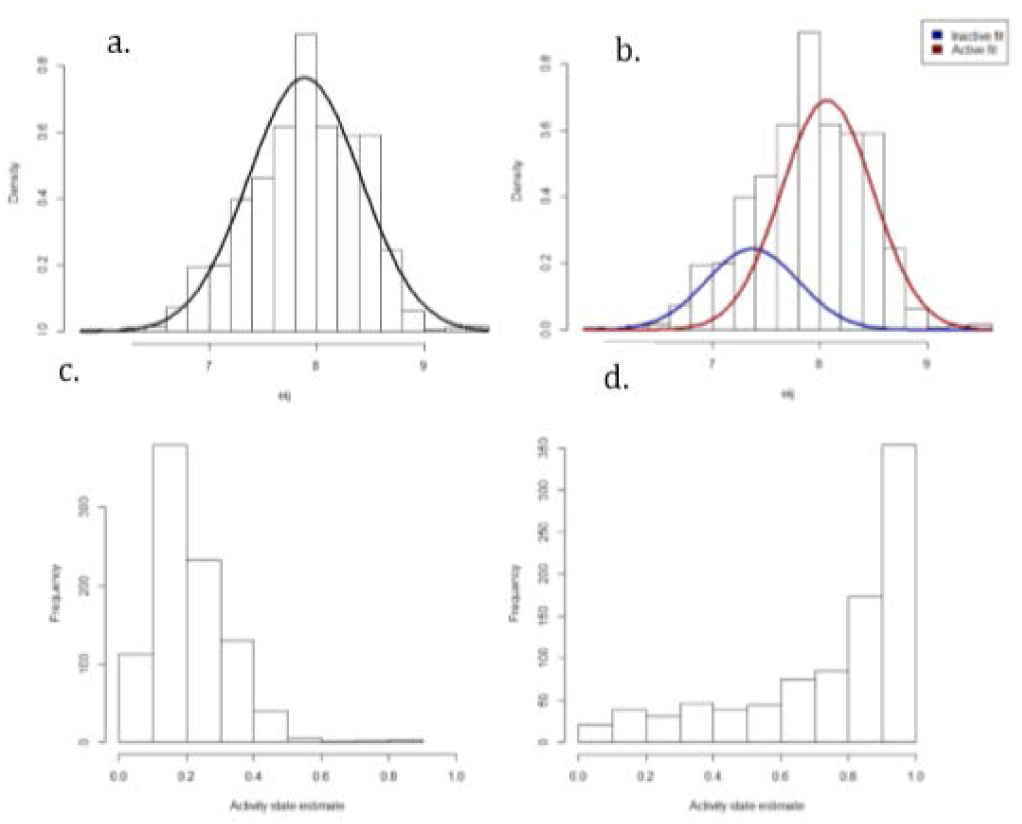
Top: expression data for *E. coli* gene *ymjA* across 907 unique experiments, with its 1-component (a) and its 2-component (b) fit overlaid. Bottom: activity state estimates (*a_ij_*) for *ymjA* in the 907 experiments, according to the models presented in (a) (corresponding to c) and (b) (corresponding to d), respectively. The distribution of activity state estimates is quite different based on which method is used, despite only having a difference of 0.7 in BIC (representing strong uncertainty in which model (a) or (b) is correct). Graph (c) suggests that *ymjA* is inactive in most experiments, whereas graph (d) suggests that *ymjA* is active in most experiments.

**Figure 2.**
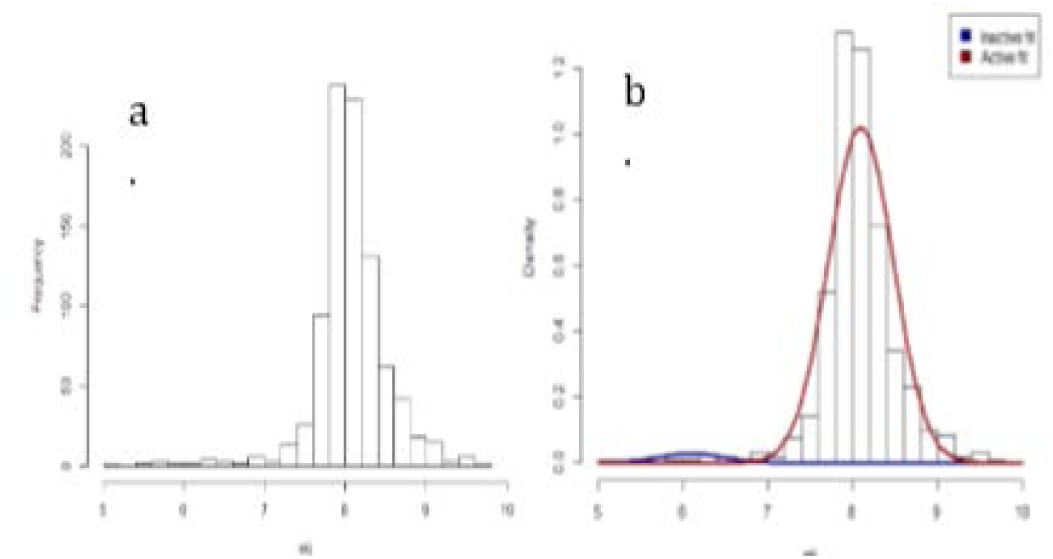
(a) Expression data for E. coli gene *rhaB* across 907 diverse experiments; (b) expression data for gene *rhaB* with its 2-component fit from *MultiMM* overlaid. The current *MultiMM* method fits a 2-component model to this gene with the ‘inactive’ component of the model fit to a small number of very low expression values. Thus, the current *MultiMM* method suggests that *rhaB* is active in most experiments

First, we let *C_i_* represent the posterior probability that gene *i* is truly changing state. We note that the current *MultiMM* approach assumes that *C_i_* ∊ {0,1}, indicating complete confidence that the gene is not-changing-state (*C_i_* =0) or is changing-state (*C_i_* =1) respectively and using BIC criterion described above (12 point lower BIC per Raftery [7]). Next, we define the “Normalized Bayes Factor” of a set of *p (p* ≥1) co-regulated genes as

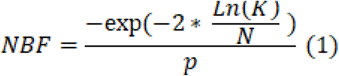
where *Ln*(*K*) is the natural log of the Bayes Factor, *K*, for the fit of the 1-component Gaussian distribution to the expression data over the 2-component Gaussian distribution as calculated by the *R* package *Mclust* [8] giving the difference in log-likelihood between the two models. Intuitively, -2 times the natural log of the Bayes factor, *K*, is a common measure of statistical evidence, and scaling by *N* (the number of experiments) and *p* provides a standardized statistic across co-regulated set size and number of experiments. We note that the highest observed value of *NBF* on our real set of 907 expression arrays for over 4000 genes is approximately 0, and the lowest observed value is -0.913, which was observed on the three genes in the *araBAD* operon. Previously, it has been noted that strong biological and statistical evidence confirms the *araBAD* genes are changing states over these 907 expression arrays [7]. We propose a piecewise linear mapping from *NBF* to *C*,

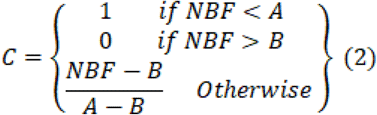
where *A* and *B* are both constants such that *A<B*. In our exploration of different cutoff choices (detailed results not shown), we observe that for genes that do not change states (true 1-component multivariate Gaussian distributions), *Ln*(*K*) appears independent of *N* and rarely falls below *-2p-3*; hence, reasonable choices for A and B are:

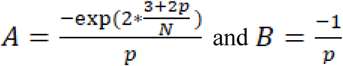

We now briefly describe how *C* is used in the generation of gene activity state estimates. In short, this approach estimates gene activity values assuming the gene (or set of co-regulated genes) is from a 1-copmonent mixture and, separately, from a 2-component mixture and then combines the estimates reflecting confidence in the estimates.

*Step 1*. For each *a priori* specified set of co-regulated genes *H*, generate gene activity state estimates, *a(2)_ij_*, using the *MultiMM* approach and assuming that the set is changing states.

*Step 2*. For each gene, *i* in the set, find parameters μ_i_ and σ_i_ describing the best-fit 1-component Gaussian distribution using *Mclust*. Then, looking at the full list of genes for which transciptomics data is available, make a list *L* (of length *n*) of all genes *g* where σ_i_ ∊ σ_g_ ± 0.1 and either μ_i_ ∊ μ_0,g_ ±0.1 or μ_i_ ∊ μ_1,g_ ±0.1.

*Step 3*. Generate *n* sets of gene activity state estimates for *i*, one for each gene in *L*, as if *i’*s expression data came from the other gene’s best-fit 2-component distribution. Let these estimates be called *a(1)_ij_* for each gene *i*.

*Step 4*. For each gene *i*, activity state estimates are calculated as

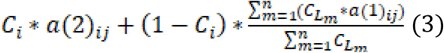

Step 5. Following the original *MultiMM* approach, for each co-regulated gene set and each experimental condition, use the average activity state estimates for each gene in the set as the final activity state estimate for each of the set’s genes.

Note that this method reduces to the standard *MultiMM* approach f *C_i_* = 1 or *C_i_* = 0.

#### ii. Incorporating uncertainty into a priori sets of co-regulated genes

The *MultiMM* method fits a *k*-dimensional multivariate Gaussian mixture model on expression data for a set of *k* genes that are indicated to be co-regulated; a model which assumes that, in any given condition *j*, all *k* genes in the set will either be active or inactive. In cases where *k=1*, a univariate Gaussian mixture model is fit to the data. The limitation of this approach is that there is no consideration of uncertainty in the *a priori* specified set of *k* genes_[9]. To allow for this uncertainty, we propose *ConfMM*, which takes as input a single number *C_h_*, ranging from 0 to 1 for each set of co-regulated genes, representing the posterior probability that the genes in *h* are in fact co-regulated. For a set of purportedly co-regulated genes, *h, ConfMM* first conducts the *MultiMM* approach on each gene in the set separately, producing gene activity estimates *U_ij_* for each gene *i* and experimental condition *j*, and then conducts *MultiMM* on the entire set of genes as a unit, producing gene activity estimates *M_ij_*. For each gene *i* and experimental condition *j, ConfMM*’s output gene activity estimates are then given by 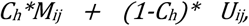 where *H* is the co-regulated set containing *i* Similar to the approach for modeling uncertainty in whether or not a gene is changing states (section i), here we take a weighted average of the gene activity estimates to capture the uncertainty quantified by *C_h_*.

#### iii. Outlier Handling

Gaussian mixture modelling can be sensitive to extreme values in the data caused by measurement and other errors, ultimately leading to biased estimates of gene activity status. The current *MultiMM* approach takes the view that outliers are biologically meaningful and not due to error. We propose two potential methods for mitigating the influence of outliers on downstream gene activity state estimates.

-*SD(d)*: This approach first calculates the mean 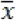 and the standard deviation *S* of the expression data for each gene, and then removes any observations greater than 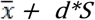 or less than 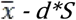.

*-Wins(d):* This approach, called “winsorization” [10], is similar to *SD(d)*, but does not actually remove any observations. Instead, any observations greater than 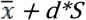 are imputed as 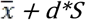. and likewise any observations less than 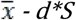 are imputed as 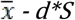. To evaluate winsorization, we consider *d=* 4 and *d= 5*.

### B. Real Data Sets

We use genome-wide expression data comprising 4329 *E. coli* genes from 907 different microarray data sets in a variety of diverse conditions. Raw data from Affymetrix CEL files were normalized using RMA [11] and these data were placed in the M3D data repository [12]. Further details of data processing are described in [13], [14]. *E. coli* operon predictions for 2648 operons (co-regulated gene sets), including 1895 single gene operons, were obtained from Microbes Online [9].

### C. Simulated data sets

For some analyses, we used simulated data with characteristics based on the real set of 4329 *E. coli* genes across 907 different experiments, with co-regulated sets of genes based on prior predictions [9]. To mirror real expression data as closely as possible, we first screened the simulated data and dropped all co-regulated sets of genes (operons) for which the 2-component model did not yield the highest BIC. A random selection of the remaining operons was chosen to be single component in the simulated data with each of the single component operons being always active or inactive with equal likelihood. In order to demonstrate our method’s performance in the face of uncertain operon classifications, we generated a second simulated data set. We simulated 100, 3-gene sets over 907 experiments, where some proportion *p* of the 100 gene sets were actually co-regulated, and the other *100(1-p)* gene sets were actually generated as independent genes. In each test we then assigned confidence *p* to each of the 100 gene sets. We will refer to this simulated data set as the *Co-regulated Confidence* data.

### D. Alternate gene activity state estimation methods

Following our earlier work [5], in this paper we consider a variety of methods for inferring gene activity states: *MT* (*Median Threshold*: all genes in an experiment above the median are deemed active and, those below, inactive), *TT (Trichotomous Threshold*: all genes below the 40^th^ percentile are deemed inactive, above the 60^th^ percentile are active, otherwise gene activity is deemed ‘uncertain’), and *RB (Rank-based*: the activity state estimate is the percentile rank within the experiment). The *MT* approach can be found in [15]. *TT* is an extension following GIMME which allows for an “uncertain” classification as in [15], and *RB* is a further extension in the spirit of GIM3E [16]. Further discussion of such methods can be found elsewhere [5]. We also consider alternatives to the *MultiMM* method: *MultiMM(12)* is the standard *MultiMM* approach using a difference of 12 in BICs to determine if a gene is changing state; *MultiMM(0)* is a variation where no preference is given when selecting based on BIC. *UniMM* is a univariate version of *MultiMM* that simply discards all co-regulation data and processes each gene separately. Finally, *MultiMM(NBF)* is the version of *MultiMM* proposed earlier (i.) which uses *NBF* to estimate *C*.

### E. Validation and statistical analysis

For simulated data, true gene activity states are known by nature of the simulation. For real data, predictions of gene activity were obtained using FVA [17] on the *E. coli* iJO1366 metabolic model [18]. For the purposes of evaluating the performance of the methods we use the “mean square alignment”, defined as

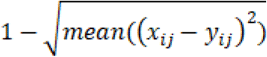
where *x* and *y* represent sets (usually matrices) of probabilities, and the mean is taken over all *i* and *j* Usually, *x* will be a set of gene activity state estimates from the method being evaluated, and *y* will be either the true gene activity states (on simulated data) or the FVA predictions (on real data). Higher mean square alignment indicates improved agreement (that is, less difference) between the sets.

## III. Results

### A. Hedging to improve inference about whether a gene is changing activity states

Figure 3 shows the mean square alignment between various methods’ *C* estimates and the true classifications, on the *Sim-Uniform E. coli* data. This shows that *UniMM(NBF)* and *MultiMM(NBF)* provide the best overall performance (measured as mean square alignment between the *C* values and the true classifications) among univariate and multivariate procedures, respectively.

**Figure 3.**
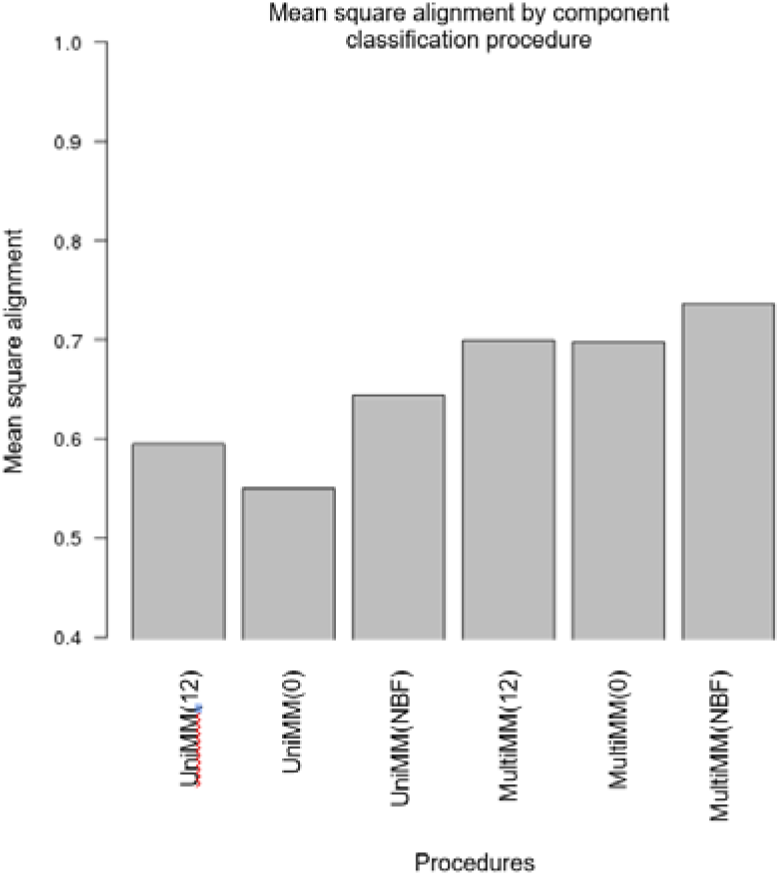
Performance of various classification procedures on Sim-Uniform E. coli data.

### B. Co-regulated gene set confidence levels

Figure 4 illustrates the performance of the *ConfMM* method on the *Co-regulated Confidence* simulated data described above. In particular, the performance of the three methods are evaluated as the proportion of true co-regulated genes in the set (*p*) varies. Notably, and by design, *ConfMM* is as good as, or better than, both *UniMM* and *MultiMM* in every case.

**Figure 4.**
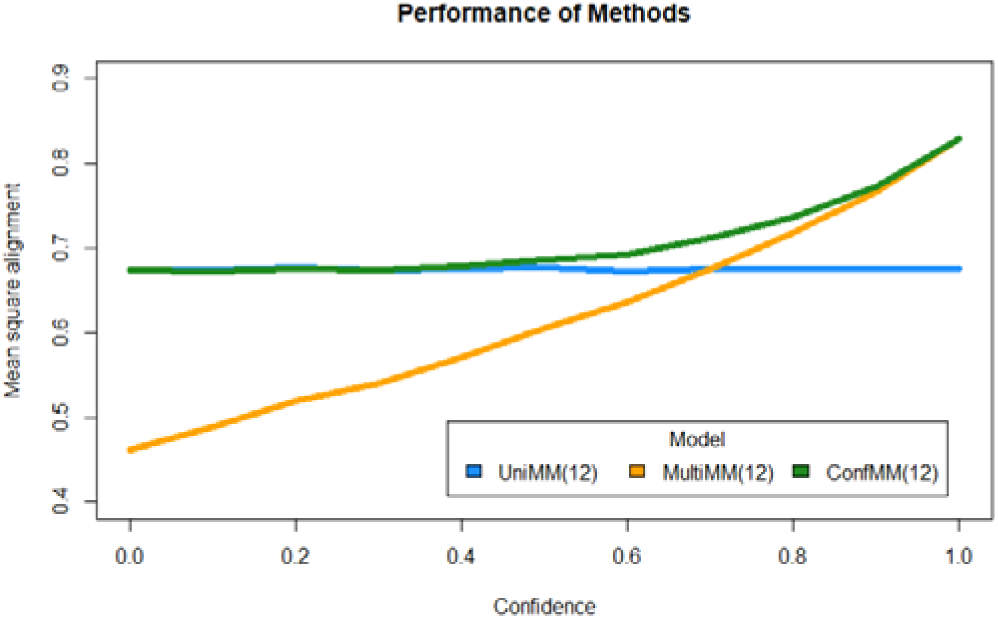
Performance of UniMM, MultiMM and ConfMM with varying levels of operon certainty.

### C. Outlier Handling

Performance of each of the gene activity state estimation methods with each of the outlier-handling approaches, as measured by mean square alignment with the FVA predictions on the real data, is given in Figure 5. We can see an illustration of the effects of outlier-handling by returning to *rhaB*, the gene from Figure 2. The raw expression data for *rhaB* was shown in Figure 2(a); the expression data after *SD(4)* or *Wins(4)* preprocessing is shown in Figure 6(a) and 6(b) respectively. The series of unusually low observations in Figure 2(a) were partially removed (by *SD(4))* or made less extreme (by *Wins(4))*. In Figure 2(b) we saw that in the raw expression data for *rhaB* (no outlier handling), the series of unusually low observations was interpreted as an ‘inactive’ cluster, with the entire rest of the observations then interpreted as ‘active’; this results in the gene activity estimates shown in Figure 6(c) whereby *rhaB* is very confidently identified as ‘active’ in the vast majority of experiments. After *SD(4)* preprocessing, the expression data for *rhaB* (shown in Figure 6(a)) was classified by *MultiMM(0)* as not-changing-state, whereas the original expression data for *rhaB* (shown in Figure 2(a)) had been classified by those methods as changing-state, leading to the interpretation explained earlier. This leads to substantially different gene activity estimates, shown in Figure 6(d). If *Wins(4)* preprocessing is used rather than *SD(4)*, a third interpretation results (see Figure 6(e)) in which rhaB is classified changing-state, but not due to the outliers on the left; instead, the overall right-skewness of the expression data is interpreted as a large ‘inactive’ cluster and a small ‘active’ cluster, producing the estimates shown in Figure 6(f). The outlier handling yields a more biologically meaningful result as rhamnose is rarely present in the 907 experiments considered here.

**Figure 5.**
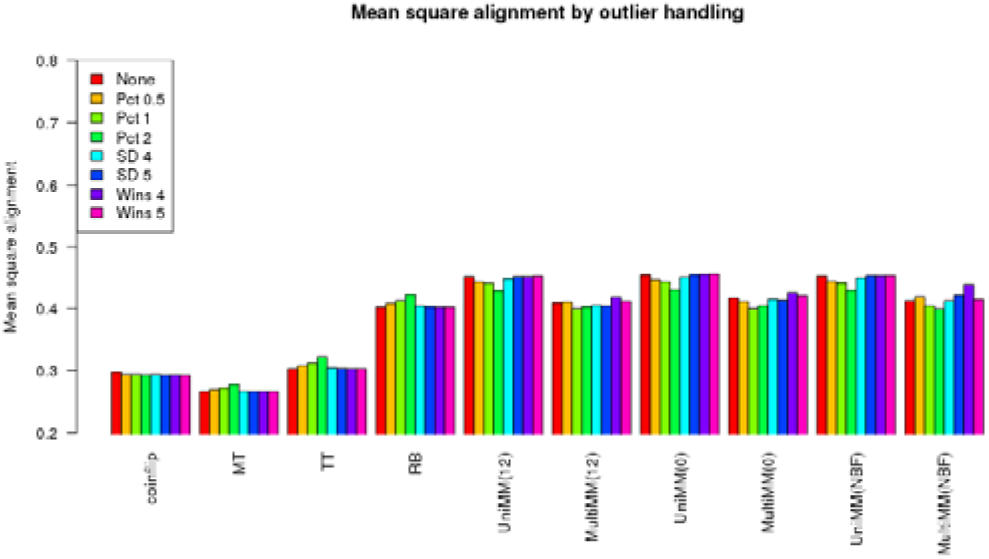
Performance of various outlier-handling methods on real data. Coin flip is a random choice as to whether the gene is active (50% chance) or inactive (50% chance). In general, outlier handling improves mean square alignment with SD and Wins methods performing relatively similarly across methods.

**Figure 6.**
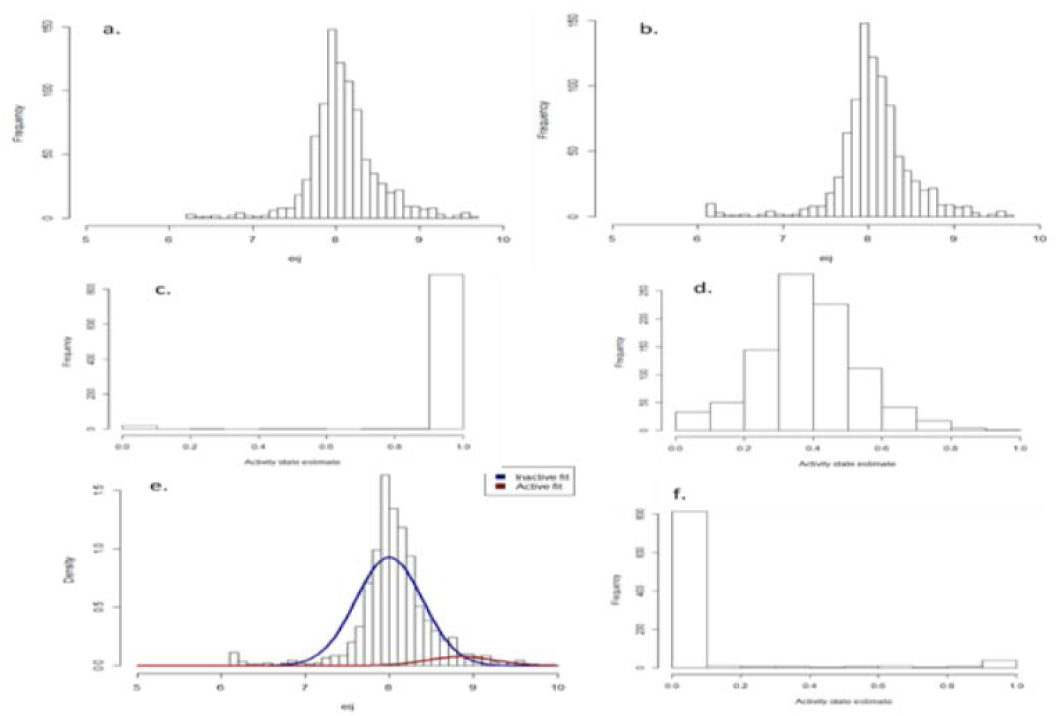
(a) and (b) Expression data for gene rhaB following *SD(4)* and *Wins(4)* preprocessing respectively; compare to Figure 2(a). (c) and (d) Activity state estimates for rhaB in the 907 experiments according to *MultiMM(0)*, with no preprocessing or *SD(4)* preprocessing respectively. (e) and (f) *MultiMM(0) Wins(4)* interpretation: 2-component fit and generated activity state estimates.

## IV. Discussion

In this manuscript we have addressed three limitations of the recently published *MultiMM* method for inferring gene activity states from genome-wide transcriptomics data: (1) accounting for uncertainty in initial inference about whether a gene is changing states, (2) uncertainty in whether a set of genes is co-regulated and (3) robustness to extreme gene expression values (outlier handling). We demonstrated that on both real and simulated data the new method performed better compared to the existing *MultiMM* method.

The Bayesian modeling framework that we present in this manuscript provides a flexible and adaptable approach to infer gene activity. Thus, as additional sources of biological information are obtained, they can be easily integrated into the framework. Next steps include leveraging both empirical estimates of gene co-regulation (e.g., correlation estimates from the set of expression data) and more precise information about regulatory relationships. One such relationship comes from the Transcription Regulatory Network (TRN). A full integration of TRN information would require the integration of the relationships themselves and explicit incorporation of TRN uncertainty into the Bayesian framework of the *MultiMM* model.

Additional validation of these method refinements and the original method are still necessary on organisms beyond *E. coli*. Furthermore, many additional refinements are possible including the incorporation of additional biological information into the Bayesian model (e.g., cross-species gene orthology, metabolic pathway information, etc.). Ultimately, these gene activity estimates can be used in multiple downstream applications including gene set analysis [6] and metabolic flux modeling, among others. Software for the methods illustrated here is available as supplemental files to this manuscript and found here: http://www.dordt.edu/statgen.

## Acknowledgments

This work is supported by National Science Foundation Grant MCB-1330734. We gratefully acknowledge the use of the Silicon Mechanics grant funded computer cluster on the campus of Dordt College for computations. The funders had no role in study design, data collection and analysis, decision to publish or preparation of the manuscript.

